# Splice-switching of the insulin receptor in rhabdomyosarcoma: Rescuing the *IR-B* isoform for better treatment options

**DOI:** 10.1101/2020.10.02.324053

**Authors:** Safiya Khurshid, Matias Montes, Daniel F. Comiskey, Brianne Shane, Eleftheria Matsa, Chelsea Brown, Hemant Kumar Bid, Ruoning Wang, Peter Houghton, Frank Rigo, Dawn Chandler

## Abstract

Rhabdomyosarcoma (RMS) is an aggressive pediatric tumor with poor prognosis for metastasis and recurrent disease. Large scale sequencing endeavors demonstrate that RMS tumors have limited mutations and a dearth of driver mutations that are precisely targetable. However, IGF2 signaling is known to be grossly altered in RMS. The IGF2 signalling molecule binds both its innate IGF1 receptor as well as the insulin-receptor-variant-A (*IR-A*) with high affinity. Mitogenic and proliferative signalling via the canonical IGF2 pathway is therefore augmented by *IR-A*. The insulin receptor (*IR*) which is a transmembrane tyrosine-kinase receptor exists in two alternatively spliced isoforms, *IR-A* and *IR-B*. In this study, we show that RMS patients express increased *IR-A* compared to control tissues that express predominantly the *IR-B* isoform. We also found that *Hif1a* is significantly increased in RMS tumors, portraying their hypoxic phenotype. Furthermore, the alternative-splicing of *IR* adapts to produce more *IR-A* in response to hypoxic stress. Upon examining the pre-mRNA structure of the gene, we identified a hypoxia-responsive-element, which is also the binding site for the RNA-binding protein *CUG-BP1*. We designed Splice-Switching-Oligonucleotides (SSO) against this binding site to decrease the levels of *IR-A* in RMS cell-lines and consequently rescue the *IR-B* expression levels. SSO treatment resulted in significant reductions in proliferation, migration and angiogenesis. Our data show promising insight into how impeding the IGF-2 pathway by reducing *IR-A* expression mitigates tumor growth. Our data reveal that RMS tumors use *IR* alternative-splicing as yet another survival strategy which can be exploited as therapeutic intervention in conjunction with already established anti-IGF-1 receptor therapies.

## INTRODUCTION

Rhabdomyosarcoma (RMS) is the most common soft tissue sarcoma of childhood, with an incidence of 4.5 cases per million children/adolescents per year in the United States ^1,2^. It is divided in two main histological variants: embryonal (ERMS, 60–70% of all RMS cases) and alveolar (ARMS, approximately 30% of all RMS cases) RMS. Despite improvement in the treatment, there is substantially poor prognosis for relapse and metastasis patients. Overexpression of IGF2 has been found in RMS tumors of both histological subtypes which makes the IGF signaling pathway one of the most frequently deregulated signaling pathways in RMS ^3–5^. Therefore, it is imperative to find novel ways to impede the IGF pathway and invent novel therapeutic strategies for the treatment of this cancer.

Alternative splicing of the insulin receptor generates two isoforms: the full-length *IR-B* and exon 11-skipped *IR-A* isoform. *IR-B* is highly expressed in adult, insulin-responsive tissues (skeletal muscle, liver and adipose tissue) and promotes the metabolic effects of insulin. *IR-A*, on the other hand, is predominately expressed in embryonic tissue and signals proliferative effects^6,7^. The insulin receptor (IR) is a trans-membrane protein from the family of tyrosine kinase receptors that is activated by insulin, IGF-1 and IGF-2, and has an isoform-specific affinity towards each ligand. IR-A and IR-B have similar affinities for native insulin, but differ substantially in their affinities for IGF-1 and IGF-2^8^. The increased affinity of *IR-A* towards IGF2 orchestrates a cascade of signals involved in numerous developmental and mitogenic pathways^9^. A study performed using the insulin receptor minigene, which consists of exons 10,11 and 12, has shown *SRSF3* and *CUGBP-1* respectively as the potential negative and positive regulators that help mediate the insulin receptor-splicing event ^10^. However, not much is known about the interplay of these RNA-binding proteins and their response to stress. Over-activation of this pathway by both insulin and IGF-2 is common in cancer cells, particularly in undifferentiated/stem-like cells, and may represent an important factor of resistance to various anti-cancer drugs^11,12,8^. Dysregulated expression of *IR-A*, which tightly binds IGF-2, contributes to these mechanisms.

In this work we show that pediatric RMS patient samples as well as RMS cell lines demonstrate an increased expression of *IR-A* as compared to the normal muscle tissue. Concordantly, the insulin receptor isoform expression increases in hypoxic conditions. Furthermore, we show that the splice switching oligonucleotides (SSOs) are able to reverse this splicing. This decrease in the expression of the *IR-A* isoform results in significantly reduced proliferation, migration, and angiogenesis of RMS cell lines. Overall, our data delineate a novel strategy to mitigate the tumor signaling by modulating the splicing of insulin receptor from *IR-A* to *IR-B*.

## RESULTS AND DISCUSSION

### The alternately spliced insulin receptor-A isoform is prevalent in RMS patient samples and cell lines

The insulin receptor gene is composed of 22 exons. Whereas, all exons are included in normal adult tissues, exon 11 is excluded during embryogenesis. Exon 11 of the *IR* gene is composed of 36 nucleotides that encode for 12 amino acids residing at the C-terminal of the alpha subunit of the protein. Though only a small change in the protein composition, deletion of these 12 amino acids results in a receptor that has increased binding affinity for the IGF-2 growth factor and is capable of responding to autocrine and paracrine signaling (Figure 1A). The expression of these isoforms is regulated during development and is altered in some breast and liver cancers^4,5,13^. RMS is characterized by high levels of IGF-2, produced in an autocrine manner ^9^. In addition to this, an increase in the expression of IGF-2R has been detected in several RMS tumors^14,15^. It is evident that IGF-1R, and its ligands play roles in proliferation and survival of tumor cells. Although there have been efforts to target and block the IGF-2 pathway by either using IGF-1R or AKT/ mTOR antibodies, only low response rates have been achieved. This can be attributed to multiple reasons, but primarily because single-agent therapies are insufficient, short-lived, and some of them display high toxicity. Additionally, this loss of efficacy might result from a compensatory signaling of *IR-A* binding to IGF2 ^16^. Based on these observations, we hypothesized that RMS tumors might display an aberrant alternative splicing of the insulin receptor pre-mRNA and sought to understand its splicing pattern in these specific tumors. To this end, we analyzed normal muscle samples and multiple RMS cell lines and performed RT-PCR for the *insulin receptor* gene. The primers flanking exon 11 amplify two products - *IR-B* and smaller exon 11 lacking *IR-A*. By calculating the ratios of these products, we found that the expression of *IR-A* is increased from an average 23.6% (*SEM 2*.*66*) in normal muscle to 80.7% (*SEM 2*.*72*) in RMS cell lines. We further performed this analysis in control muscle tissue and a cohort of 30 embryonal and alveolar rhabdomyosarcoma patient samples. In 100% of the tumor samples, we witnessed an increased expression of the *IR-A* isoform compared to the normal controls (Figure 1B, C, D). These data show that RMS tumors change the splicing pattern of insulin receptor to express more *IR-A*, which, based on the significantly higher affinity of *IR-A* for IGF2, might partly explain the increased downstream IGF-2 signaling in these patients ^9,17^.

**Figure 1:**
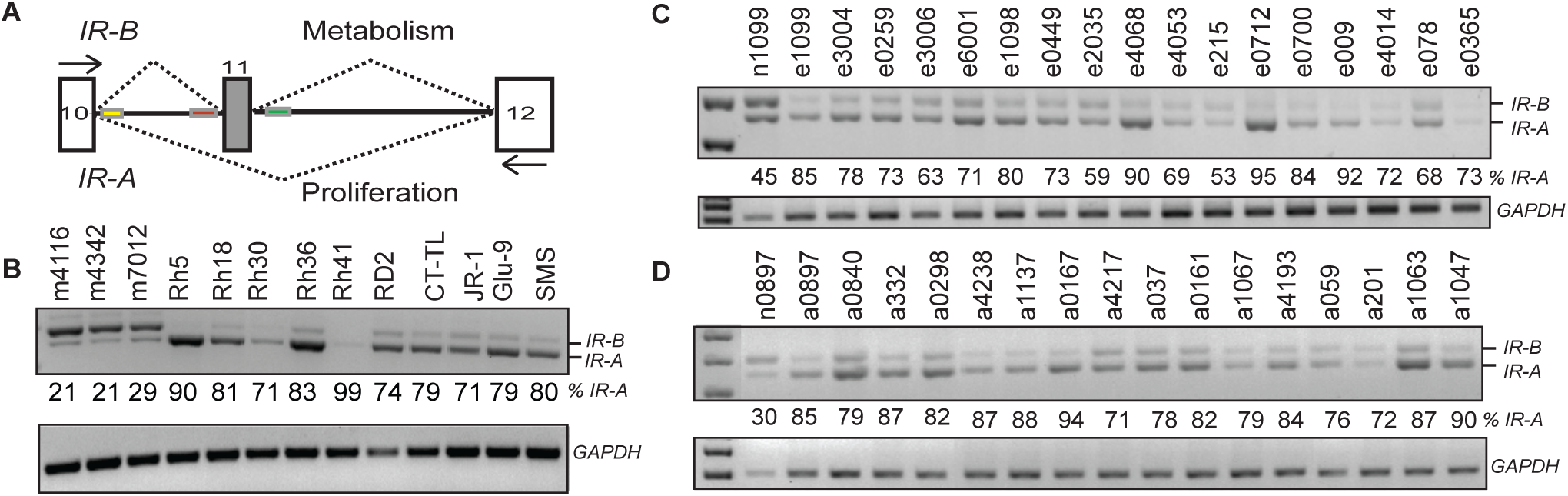
RMS cell lines and patient samples express increased levels of *IR-A*: **A**. Cartoon depicting the splicing of insulin receptor gene. **B**. RT-PCR of control muscle samples and RMS cell lines depicting the 2 insulin receptor (IR) isoforms *IR-A* and *IR-B*. Cells were cultured in the required medium, RNA was extracted and PCR was performed using the primers depicted by black arrows in A. The PCR products were run on a 1.5% agarose gel and bands were quantified. RT-PCR performed on **C**. Embryonal and **D**. Alveolar RMS samples. GAPDH is shown as control.

### Hypoxia increases the expression of *IR-A* isoform and mutation of the hypoxia response element in the minigene construct prevents it

Solid tumors are characterized by both transient and chronic hypoxia, making the ability of tumor cells to adapt to hypoxia essential for tumor progression^18^. In addition to disruption in IGF signaling pathways, hypoxia is a well-known feature of aggressive pediatric tumors^19,20,21^ including RMS ^22,23^ and correlates significantly with the patient outcome^24^. Some papers have identified the effect of hypoxia on the alternative splicing, but the targets have not been well characterized and the process appears to be cell type specific ^25,26^. In addition, hypoxia is known to interfere with the insulin signaling in various cell types^27,28^. To better understand the relevance of the tumor hypoxia in RMS, we interrogated the Schafer-Welle 56-MAS5.0-u133a data set in which 26 normal muscle samples were compared with 30 RMS patient samples (15 ARMS, 15 ERMS) and found that RMS patient samples significantly overexpress *HIF-1α* (Figure 2A) ^29^. In order to assess the changes in *IR* splicing in response to hypoxia, we utilized the HeLa S3 cells because these cells display equal expression of the two insulin receptor isoforms. Equal number of cells were seeded and maintained in either hypoxic or normal oxygen conditions as control. After 48 hours, we found that cells that underwent hypoxia treatment had significantly increased *IR-A* induction as compared to the control cells (Figure 2B). To assess hypoxia induction, we quantitated *HIF-1α* and found it to be upregulated in hypoxia-treated cells (Figure 2B). Our results indicate that hypoxic stress induces the expression of the *IR-A* isoform in cancer cells. Our findings support the idea that RMS cells induce differential splicing of the *IR* gene in order to adapt to the hypoxic microenvironment and potentially promote tumorigenesis.

**Figure 2:**
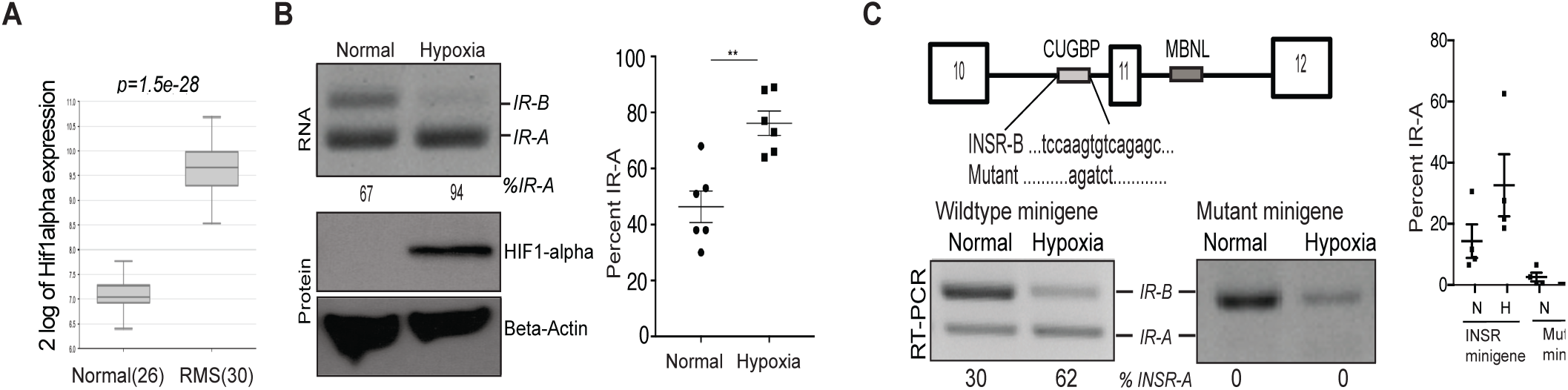
Hypoxic stress increases the expression of *IR-A* isoform: **A**. The Gene expression data for HIF-1α expression were extracted from *Schafer-Welle* (*Scafer-Welle-56-MAS5*.*0-u133a*) dataset using R2: Genomics Analysis and Visualization platform for 26 normal muscle samples and 30 RMS samples^35^. **B**. RT-PCR depicting the IR alternative splicing in HeLa S3 cells after 48 hours of normoxia or hypoxia (1% O2). Quantification of *IR-A* expression in control and hypoxic conditions; n=6, *p=0*.*0018*. **C**. Depiction of the wild-type insulin receptor (*IR*) minigene or the *CUG-BP1* binding-site mutated minigene. RT-PCR depicting expression of insulin receptor isoforms using an IR control minigene and a *CUG-BP1* binding-site mutated minigene. Quantification of C.

In order to understand the specific regulation of the *IR* splicing process, we obtained an insulin receptor minigene, which is comprised of exons 10, 11 and 12^10^. We transfected RMS cells with the *IR* minigene and subjected them to hypoxic conditions. We found that skipping of exon 11 is increased after hypoxia treatment showing that the minigene retains the hypoxia-induced alternative splicing response similar to the endogenous gene. To interrogate the role of *CUG-BP1*, a known regulator of the *IR* splicing event^10^, we deleted its binding site in the minigene and found that the *IR-A* isoform could no longer be induced, as was the case with the wild-type construct (Figure 2C) showing that this binding site in the intron-10 contains the hypoxia responsive element, which is essential for this splicing function.

### Splice switching oligonucleotide (SSO) treatment restores *IR* splicing to *IR-B*

Based on the information of hypoxia responsive element (Fig 2), as well as the CUG-BP1 binding sites in the insulin receptor pre-mRNA, we designed splice-switching oligonucleotides (SSOs) to increase the inclusion of exon 11 by masking the negative regulatory element within the *IR* pre-mRNA. We also designed SSOs to exclude exon 11 by masking a downstream positive regulatory element within the *IR* pre-mRNA that was previously identified^10^. The SSO methodology allows the *IR* gene to be expressed only from the endogenous locus and at levels expressed by normal cells. The SSOs have been used successfully in the past both to decrease inclusion of exons (in the muscular dystrophy-causing gene *DMD*) and to increase inclusion of exons (exon 7 of the spinal muscular atrophy [SMA] causing gene *SMN*) ^30^. In order to identify the *CUG-BP1*-binding region, which we showed to be important for hypoxia induced splicing change of the *IR* transcript, we screened nucleotides on both sides of exon 11. We co-transfected HeLa S3 cells with an *IR* minigene as well as the designated SSO. As a negative control we used an SSO (SSO23) that we previously identified to cause increased exclusion of exon 11, thereby increasing *IR-A* (Jung & Chandler unpublished). SSO23 blocks the binding site for the *MBNL1* protein, a positive regulator of *IR* splicing, and successfully targets a nucleotide element in *IR* intron 11 to increase the levels of *IR-A* (Figure 3A).

**Figure 3:**
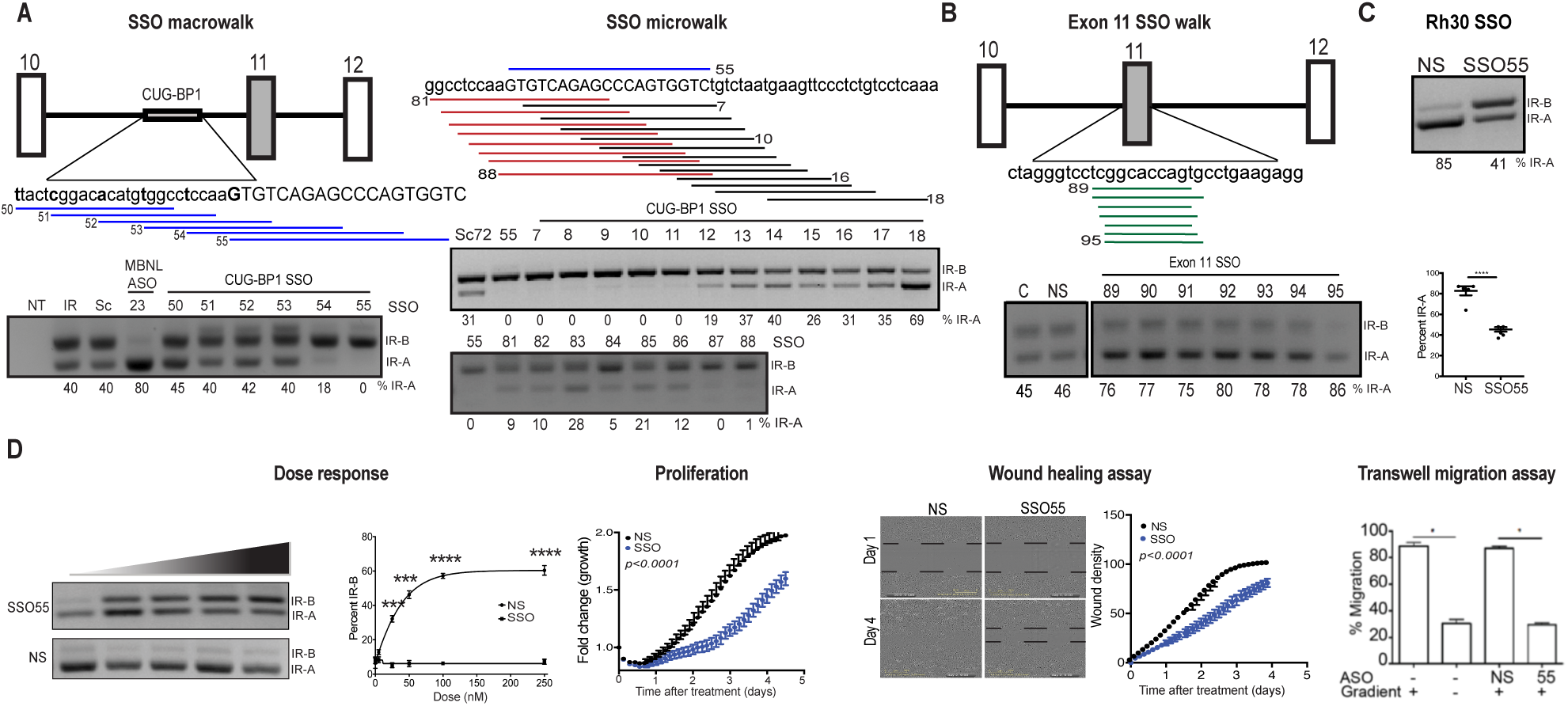
Splice switching oligonucleotide (SSO) treatment restores *IR* splicing to *IR-B*. **A**. Representation of the insulin receptor minigene. The *CUG-BP1* binding site is shown on the *IR* pre-mRNA. The blue lines represent the SSOs used for the macro-walk and the red and black lines represent the SSOs used for the micro-walk. RT-PCRs depicting the IR isoforms in the presence of control SSOs or SSOs that target either the CUG-BP1 binding site are shown below the sequences. **B**. Represenation of exon 11 SSOs (green lines). RT-PCR using SSOs on the second *CUG-BP1* binding site in exon 11 is shown below. **C**. Non-specific (NS) and SSO55 were transfected endogenously into Rh30 cells and RT-PCR for *IR* was performed after 24 hours. The graph below RT-PCR shows the quantification from n=5 experiments. **D**. Dose Response: 500000 cells were seeded, transfected with different quantities (0-20nm) of non-specific (NS) and SSO55. RNA was extracted after 24 hours and RT-PCR for insulin receptor was performed. Dose response curve of SSOs showing the IR isoforms and their quantification in Rh30 cells. Proliferation: Rh30 cells were seeded, transfected with either NS or SSO55 and subjected to proliferation assay using the Incucyte® software. Wound healing assay: Rh30 cells were seeded in a 96-well plate and transfected with either non-specific (NS) or SSO55. After 24 hours, the Incucyte® wound maker was used to make the wound and the migration of cells was measured using the Incucyte® platform. The wound density numbers are quantified. Statistics are calculated using 2-way ANOVA. Transwell migration assay: Rh30 cells were transfected with NS or SSO55 and placed in the top chamber of a dual-chamber Boyden assay and exposed to an FBS gradient or not. Untransfected Rh30 cells were used as control. Percent migration was quantified. n=4 for each group, asterisk indicates *p=0*.*05* in one-way ANOVA with Holm-Sidak post-hoc analysis.

In order to discern whether we can use the SSOs to restore the inclusion of *IR* exon 11, we performed a ‘macro-walk’ using six overlapping SSOs (SSO50-SSO55), represented by blue lines, spanning the entire region in 5-nt increments to target a 40-50 nucleotide region containing the *CUG-BP1* binding site in intron 10 of *IR*. Out of the five SSOs, the last two were capable of inducing *IR-B* expression from 60 to 82%, with SSO55 reaching the maximum effect and increasing the *IR-B* to 100% (Figure 3A). In order to pinpoint the most effective SSO targeting this sequence element, we designed a ‘SSO micro-walk’ around the region of SSO55 by using 20 additional consecutive SSOs, represented by red and black lines (SSO micro-walk) at 2-nt resolution to cover the entire 40-nucleotide region (Figure 3A). Based on the percent expression of *IR-B* in these cells, we were able to determine that SSOs (55, 87 and 88) were highly effective in mediating this splicing change to restore the expression of *IR-B*. We tried to also use consecutive SSOs to block the 30-nucleotide element in exon 11, but were unable to target this second *CUG-BP1* binding site, most likely due to the small exon size and interference with the U1 and U2 snRNPs that recognize the splice sites flanking the exon or the interaction with other splice regulatory elements within the exon (Figure 3B).

As shown in Figure 1, RMS cell lines predominantly express *IR-A*. In order to define SSOs as biologically relevant agents in RMS cell line Rh30, we transfected control and SSO55 in these cells. SSO55 successfully decreased the *IR-A* isoform from 85 to 41% in an endogenous gene setting (Figure 3C). Furthermore, in order to determine an optimal dose of SSO for endogenous transfection of RMS cell lines Rh30, Rh18 and RD cells, we performed a dose response assay ranging from 0 nM to 250 nM of SSO. RT-PCR of these cells depicted a 0 to 60% increase in *IR-B* isoform and therefore restored the splicing in these cells in an endogenous setting (Figure 3D, Supplementary Figure 1).

Our data outlines a novel approach where we can use a SSO compound to reverse splicing in RMS cell lines. In addition to this our data potentially links the RNA binding protein *CUG-BP* with insulin receptor and this interplay between the two could further be exploited in therapeutic and mechanistic directions and a recent paper documented this interaction in breast cancer cells^13^.

### SSO treatment decreases proliferation and migration of RMS cell lines

The transfection of SSO55 in Rh30 cells effectively promoted exon 11 inclusion and decreased *IR-A* splice isoform levels. In order to assess whether the restoration of this splicing helps alleviate any cancer cell hallmarks in these cells, we performed proliferation assays on RMS cell lines that were transfected with either a control non-specific SSO (NS) or SSO55, specific for *IR-A* reduction. We found that there was a significant decrease in proliferation of cells transfected with SSO55 as compared to the control cells (*p<*0.0001) (Figure 3D). Similarly, other RMS cell lines like Rh18 (*p<*0.001) and RD (*p<*0.01) showed a significant decrease in proliferation (Supplementary Figure 1A). In addition to proliferation, another hallmark of cancer cells is increased migration. To determine whether the splicing shift towards the *IR-B* isoform also inhibits migration, we performed scratch-wound assays in Rh30 cells transfected with SSO55 or NS-SSO. We found that while the control cells migrated faster and closed the gap in 90 hours, the cells transfected with SSO55 showed a decreased migration rate and failed to close the gap within that time (Figure 3D). Furthermore, we performed quantitative migration assays and found that SSO55 decreased the migration of Rh30 cells by 6-fold (Figure 3D). Our data shows that shifting the splicing of the insulin receptor back to *IR-B* isoform inhibits proliferation and migration of RMS cells, two important attributes of cancer.

### SSO treatment diminishes tube formation and angiogenesis in RMS cells

IGF-1R blocking monoclonal antibodies showed poor response rates and largely failed in the late phase clinical trials. The failure can be attributed to the increased toxicity levels as well as hyper-insulinemia. Also, if RMS tumors have an increased *IR-A* as an alternative conduit for the downstream expression of IGF signaling, the blocking of IGF-1R is not very helpful. Related studies from the pediatric preclinical testing program (PPTP) found that the IGF-1R inhibitor shows anti-angiogenic effects^31,32^. In order to check whether angiogenesis is affected in the cells transfected with SSO55, we performed a proteome profiler array wherein we used a nitrocellulose membrane with immobilized antibodies and incubated it with biotinylated lysates from Rh30 cells that are transfected with non-specific SSO or SSO55 in the presence or absence of an IGF-1R antibody. The proteins were detected using streptavidin-HRP detection method. Our data shows that MMP9, a matrix metallopeptidase previously shown to be involved in tumor vascularization and metastasis ^33,34^ is decreased 8-fold in the presence of SSO55 and reduced to near undetectable levels in the presence of both SSO55 and IGF-1R-blocking antibody. The decrease in MMP9 levels is confirmed by the Western blot (Figure 4B, C). MMP9 has been shown to positively regulate the insulin receptor survival pathway^35^ and therefore its abrogation with the SSO works favorably to inhibit the growth of cancer cells. No other protein on the proteome array showed any changes either treatment with IR antibody or SSO alone. However, another 6 out of 55 proteins represented showed a significant decrease in the presence of both SSO55 and IGF-1R antibody and these included the major angiogenic modulators such as *VEGF, TSP-1, TGFb1*, all known to be acting downstream of *IGF-1R* (Figure 4B).

**Fig 4:**
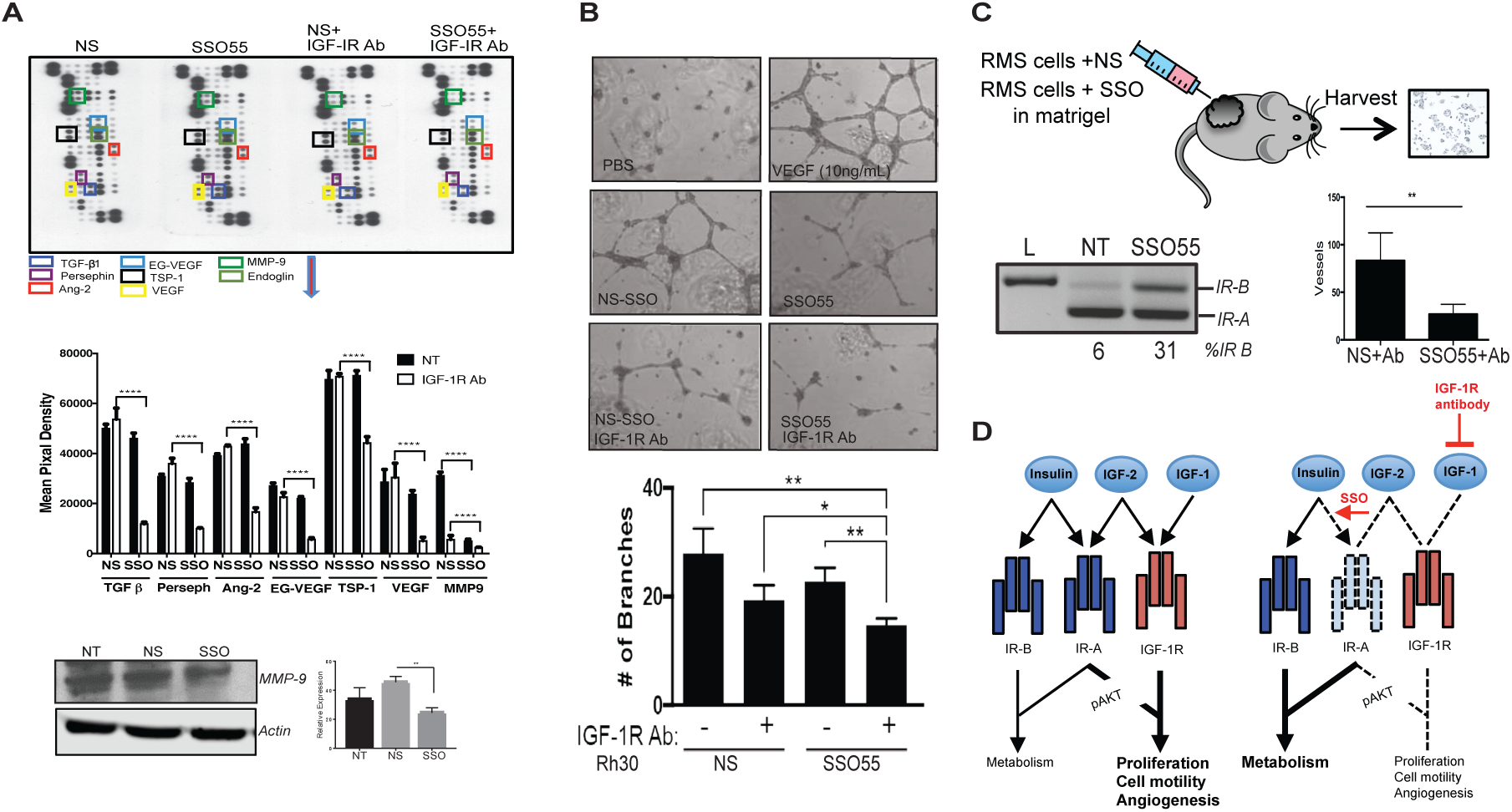
SSO treatment diminishes tube formation and angiogenesis in RMS cells: **A**. A protein profiler array was used to assess the expression of angiogenic factors and the expression was quantified. Quantification for n=2, statistics were done using 2-way ANOVA ****=*p<0*.*0001*. Rh30 cells were treated with NS and SSO55 compounds and lysed for Western blot analysis. MMP9 and Actin are shown. Quantification of MMP9 expression, n=3, Mann Whitney test. **B**. Human Umbilical Vein endothelial cells (HUVEC) were incubated with either PBS or media from Rh30 cells treated with either NS or SSO55 alone or in combination with IGF-1R antibody. The vessels/branch formation was measured and quantified using CD31 antibody staining. **C**. Mice were injected with 10^6^ Rh30 cells either transfected with NS or SSO55 and the tumors are extracted after 5 days for RT-PCR analysis and after 7 days for CD34 staining. The tumors were formalin fixed and paraffin embedded and slides were stained for CD34 and quantification. Quantification of the CD34 staining from NS and SSO55-treated grafts was done in a blinded manner, n=10 mice each condition. The RT-PCR shows the splicing changes in the injected cells in the presence of SSO55. **D**. Model depicting how insulin and IGF2 bind to *IR-A* in tumor cells leading to proliferative signaling while as in the presence of an IGF-1R antibody and an SSO that shifts the splicing to IR-B, the proliferative signaling can be mitigated.

In order to determine whether this decrease in the angiogenic factors effects the tube formation in endothelial cells, we established 2D co-cultures of HUVECs with Rh30 cells. Briefly, human umbilical vein endothelial cells (HUVEC) were plated with extracellular matrix and conditioned media from Rh30 cells treated with either NS or SSO55 with or without an IGF-1R antibody. The formation of capillary structures called tubes was measured using an inverted microscope. Our data shows that the treatment of IGF-1R antibody alone or the SSO55 alone did not inhibit the endothelial cells to form capillary structures, however the number of branches was significantly decreased in cells treated with a combination of SSO55 and the IGF-1R blocking antibody (Figure 4A).

These data indicate that the combinatorial treatment of SSO and IGF-1R blocking antibody may act synergistically to abrogate the angiogenic signaling of the IGF pathway. It has been shown that stimulation of vascular endothelial cell proliferation and angiogenesis induced by the major angiogenic factor, VEGF, is dependent upon insulin-like growth factor (IGF) signaling^36^. Importantly, *IR-A* is produced in response to hypoxia and, in consequence, *IR* splicing appears as a critical component in the progression of pediatric sarcomas by acting as a receptor for growth of tumor cells, as well as for paracrine growth of vascular cells, and hence, angiogenesis. The features place rhabdomyosarcoma in a critically unique situation wherein it is imperative to target the IGF pathway using innovative approaches.

### SSO treatment attenuates insulin receptor signaling and reduces vessel formation in *SCID* mice

In order to determine the ability of SSO55 to interfere with angiogenesis *in vivo* we used the well-characterized Matrigel Plug Angiogenesis Assay ^37^. Briefly, we injected 10^6^ Rh30 cells embedded in Matrigel into flanks of nude mice. We used Rh30 cells that had been transfected with NS SSO or SSO55 24 hours prior to implantation. We harvested the plugs at day 7 and quantified the blood vessels in these Matrigel plugs by staining the paraffin sections for CD34 and found that the tumors with SSO55 show significantly less extravasation of blood vessels compared to the control. To confirm the activity of the SSO we isolated RNA from these plugs and subjected it to PCR using human specific primers to prevent cross-reaction with the mouse insulin receptor and found that the percentage of the *IR-B* isoform was higher in the tumors that were transfected with SSO55 (Figure 4D).

Our data shows that SSO55 can effectively shift the splicing of insulin receptor to *IR-B* isoform and consequently decrease angiogenesis in grafted tumor cells. Therefore, SSOs arise as a potential viable therapeutic strategy either alone or in combination with existing IGF2 therapies for RMS patients. Overall, the somatic mutation burden is very low in RMS tumors as compared to other tumor types. Some patients show *MYCN* or *TP53* mutations, or the formation of a fusion protein between *PAX3*/*PAX7* and *FOXO1*, but those are exceedingly rare. Therefore, in general it is difficult to treat RMS by targeting the driver mutation. One of the rare classes of mutations in ERMS tumors are mutations in the RAS pathway, nevertheless, Shern *et al*. showed that while 9/43 rhabdomyosarcomas have a *RAS* mutation, targeting the RAS pathway demonstrated no detectable anti-RMS cytotoxicity^38,39^. In addition to this, the most common mutations in RMS tumors were found to be in the receptor tyrosine kinase RAS/PIK3CA axis^38^. Because the insulin receptor signaling culminates in the activation of the PI3-kinase pathway, it is worthwhile to attenuate signaling in this pathway by targeting the *IR* splicing in combination with other treatment strategies.

## CONCLUSION

Our data unveils a new paradigm of how alternative splicing regulates cellular behavior to mediate cancer-causing changes and how SSO compounds can reinstate normal splicing and impede tumor growth. Splice switching technology has been successfully employed and is an FDA approved clinical treatment in SMA patients ^30^. Given the inefficacy as well as the toxic side effects of the chemotherapy in pediatric patients with advanced cancer, it is a crucial time to devise novel, out-of-the-box therapy options to not only more cleverly modulate the signaling pathways, but also open up a new areas of therapy platform. Taken together, our data provide insight into alternative splicing changes and the consequential aberrant downstream signaling during tumorigenesis and our SSO compounds could lead to innovative RMS therapeutics that will be translatable to the treatment of other types of cancer associated with aberrant alternative splicing.

## ACKNOWLEDGMENTS

This work was funded by NCI grant R21 R21 CA195324, an OSU Pelotonia Idea Award, and foundation grants from the Elsa U. Pardee Foundation, The Sunbeam Foundation, and Cancer Free Kids (to D.S. Chandler) as well as OSU Pelotonia fellowships (to S. Khurshid, M. Montes, and D.F. Comiskey). We would like to thank Dr. Nick Webster (UC San Diego School of Medicine) for his kind contribution of IR splicing minigene constucts. We would also like to thank the members of the Chandler lab for critical review of the manuscript.

## MATERIALS AND METHODS

### Plasmids and minigenes

The *IR* 10-11-12*s* minigene was a kind gift from Dr. Nick Webster.

### RT and PCRs

Reverse transcription (RT) reactions were carried out using 1 µg of RNA using Transcriptor RT enzyme (Catalog No. 03531287001) from Sigma Aldrich. *IR* minigene PCRs were performed as previously reported. PCRs for endogenous *IR* were performed using a primer on exon 10 5’GGCTGAAGCTGCCCTCGAG3’ and a primer on exon 12 5’GCGACTCCTTGTTCACCACC3’. This PCR generates a larger full-length product (160bp) and a smaller exon-11-skipped product (135bp). The amplicons were amplified using the following primers under the standard PCR conditions: (94°C 5′, 35 cycles of 94°C 30”, 65°C 30”, 72°C 1′, 72°C 7’).

### Western blot analysis and antibodies

Cell were lysed in NP-40 buffer and equal amounts of protein were loaded in SDS sample buffer onto a sodium dodecyl sulfate-polyacrylamide gel (SDS-PAGE), blotted onto a polyvinylidene difluoride (PVDF) membrane, and analyzed for expression of using either MMP9 (R&D Systems AF911). To detect expression of β-Actin clone AC-15 (Catalog Number A5441) from Sigma was used. Protein sizes were determined using the Precision Plus Protein Dual Color Standards marker (Catalog Number 161-0374) from Life Technologies (Carlsbad, CA, USA).

### Quantitative migration assay

Rh30 cells transfected with 100 nM SSO55 or 100 nM NS were placed in the top chamber of a dual chamber Boyden assay. Un-transfected Rh30 cells were used as a control. Cells were exposed to an FBS gradient (2% upper chamber and 15% lower chamber) or no gradient as a control (2% FBS in both chambers). Percent migration was quantified by dividing the number of cells that migrated through the porous membrane by the total amount of cells (cell amount gauged by crystal violet staining and subsequent optical density measurement at 570 nm).

### Proteome profiler microarray

The protein profiler Array^™^ Human Angiogenesis kit array kit was purchased from R&D Biosystems. Briefly, antibodies have been spotted in duplicate on nitrocellulose membranes. Cell lysates treated with NS, or SSO55, with and without IGF-1R-blocking antibody were mixed with a cocktail of biotinylated detection antibodies. The sample/antibody mixture was then incubated with the Human Angiogenesis Array. Any protein/detection antibody complex present was bound by its cognate immobilized capture antibody on the membrane. Following a wash to remove the unbound material, streptavidin-HRP and chemiluminescent detection reagents are added sequentially. Light is produced at each spot, proportional to the amount of analyte bound.

### Angiogenesis co-culture experiment

We transfected Rh30 cells with NS and SSO55. After 24 hours the cells were treated with the IGF-1R antibody. After another 24 hours the cells were transferred to the upper compartment of a transwell insert with HUVEC cells plated at the matrigel coated bottom. After 48 hours, the vascular structures (tubes) in the endothelial cells were quantified using the CD31 staining^40^.

### Matrigel plug angiogenesis assay

Rh30 cells transfected with NS or SSO55 were resuspended in matrigel and injected into flanks of nude mice. Five mice were used for each NS and SSO55 and 2 million cells were injected into each mouse flank. Matrigel tumor was excised after 7 days, cross sections taken, stained with CD34 in a blinded fashion and counter stained with Meyers stain.

### Quantification of splicing ratios

Percentages of full-length and exon-excluded products were quantitated using Image-Quant TL (Version 8.1). Significance of the results was assessed using the two-tailed Student’s *t*-test using GraphPad Prism (Version 6.0).

### Cell culture, growth and transfection conditions

RMS cell lines (Rh30, RD, SMS-CTR) cells were obtained from Peter J. Houghton. HeLa and Hela S3 cells were obtained from ATCC. All cell lines derived from human material have been verified by STR analysis. Experiments were performed within the first 10 passages of thawing cells. RMS cells were maintained in DMEM, whereas HeLa S3 cells were maintained in RMPI medium. Both were supplemented with 10% fetal bovine serum (Catalog Number SH3007103) from Thermo Fisher Scientific, 1X L-glutamine (Catalog Number MT 25-005 CI) from Corning and 1X penicillin/streptomycin (Catalog Number MT 30-001 CI) by Corning.

### SSO treatment

SSOs were kindly provided by *Ionis pharmaceuticals*. SSOs specific to insulin receptor pre-mRNA were transfected in cells with either Lipofectamine 2000 (Catalog Number 11668030) from Life Technologies.

