## Supplemental Figure 1 for "Splice-switching of the insulin receptor in rhabdomyosarcoma: Rescuing the *IR-B* isoform for better treatment options"

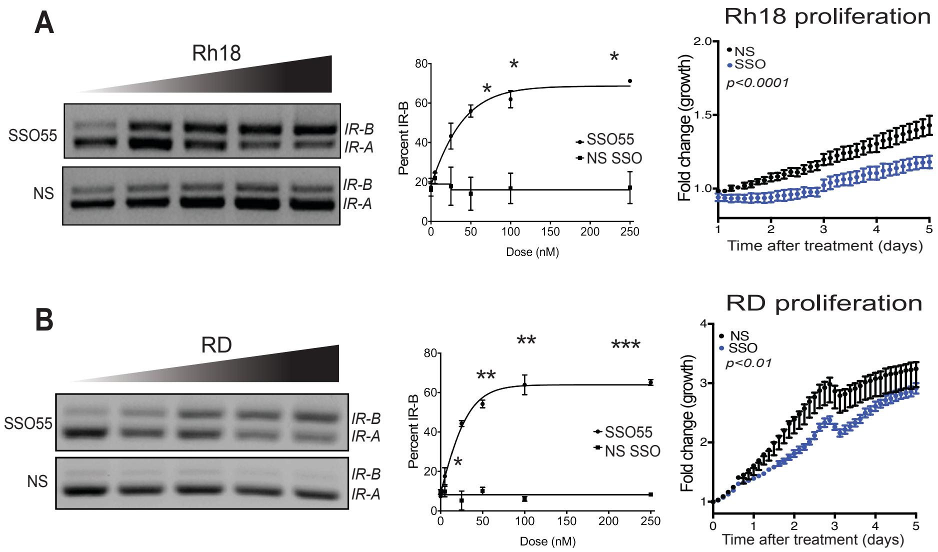


**Supplementary Fig 1. SSO treatment decreases proliferation and migration of RMS cell lines.** 500000 (**A**) Rh18 and (**B**) RD cells were seeded, transfected with different quantities (0-250 nM) of non-specific (NS) and SSO55. RNA was extracted after 24 hours and RT-PCR for insulin receptor was performed. Dose response curve of SSOs showing the IR isoforms and their quantification of (**A)** Rh18 and (**B)** RD cells. Proliferation: (**A**) Rh18 and (**B**) RD cells were seeded, transfected with either nonspecific (NS) or SSO55 and subjected to proliferation assay using the Incucyte® software.
